# The genetic consequences of dispersal and immigration in a wild great tit population

**DOI:** 10.1101/2025.06.26.660565

**Authors:** Andrea Estandía, Nilo Merino Recalde, Jon Slate, Ben C. Sheldon

**Affiliations:** Edward Grey Institute of Field Ornithology, Department of Biology, University of Oxford, United Kingdom; School of Biosciences, University of Sheffield, Sheffield, UK

**Keywords:** dispersal, immigration, local adaptation, microgeographic adaptation, gene flow, great tits, genetic diversity

## Abstract

Understanding how dispersal impacts the genetic makeup of populations is essential for predicting their responses to environmental change. Gene flow—via within-population dispersal and external immigration—shapes population health and evolutionary potential by boosting genetic diversity, but it can also counteract local adaptation. We investigate these processes in a population of great tits (*Parus major*) in Wytham Woods, United Kingdom. This system represents a large, continuous population of a vagile, widely distributed species. Using a comprehensive social pedigree alongside genomic data from two genomic datasets, one with 949 individuals genotyped at ~600,000 SNPs and another with 2,644 individuals at ~10,000 SNPs, we observe some spatial genetic structure largely driven by the spatial and temporal clustering of close kin. We quantify how temporally persistent this pattern is and find similar levels of decay of relatedness with distance across years, without a consistent genetic basis, which is frequently renewed due to high population turnover. Immigrants make up a substantial portion of the breeding population, yet are often assumed to be genetically distinct, unrelated, and outbred—assumptions that can bias population inferences. We show that immigrants are indeed outbred, as are local birds; have fewer close relatives within the population, and are less likely to be related to their neighbours than locally born birds. Despite low *F*_*ST*_ and no clear genome-wide population structure, immigrants and locals can be distinguished above chance using a Random Forest classifier trained on SNP data. Our study highlights the complex interplay between dispersal, population turnover and spatial population structure, and suggests that great tits in Wytham Woods experience substantial gene flow within the population and from immigrants, maintaining high genetic diversity and reducing the possibility of local adaptation at this spatial scale.

## Introduction

Understanding the evolutionary dynamics that shape natural populations is essential for predicting their evolutionary trajectories and responses to changing environments. Gene flow is a key driver of a population’s evolutionary potential, by maintaining standing genetic variation that natural selection can act on in the face of environmental change, counteracting the effects of genetic drift and promoting heterosis (Garant et al., 2007; Lenormand, 2002). However, it is also a homogenising force that can impede locally adaptive shifts in trait distributions. The occurrence of local adaptation depends critically on whether selection strength exceeds the effects of gene flow (Nosil & Crespi, 2004; Richardson et al., 2014), and can be severely constrained by genetic limitations and insufficient genetic variation (Orr, 2000; Slatkin, 1987).

Gene flow is more common among geographically close populations, which is why local adaptation is rarely observed at microgeographic scales, defined as the spatial scales where high gene flow is expected based on the species’ dispersal patterns (Richardson et al., 2014). However, some exceptions exist even in highly mobile organisms, such as the Californian scrub jay (*Aphelocoma californica*) (Cheek et al., 2022), the common chaffinch (*Fringilla coelebs*) (Recuerda et al., 2023), and different species of frogs (Skelly, 2004) and salamanders (Richardon & Urban, 2013), which show local adaptation to different habitat types at very fine spatial scales. These examples are mostly from islands or ponds, where immigration tends to be less frequent than on larger contiguous environments (Estandía et al., 2025). Island populations also tend to show lower dispersal than their mainland counterparts, which can encourage local adaptation and population divergence (Suárez et al., 2022; Estandía et al., 2023). On the mainland, habitat fragmentation can create a metapopulation structure, where populations are relatively isolated in small, distinct patches. In these fragmented environments, reduced gene flow between patches might allow for local adaptation. An example of this can be seen in the fragmented woodlands of the United Kingdom, where isolated populations of several bird species show population structure between woodland patches (McCollin, 1993).

Understanding the extent of gene flow across a population and the effects of external immigration is fundamental to understanding evolutionary dynamics and their evolutionary potential (Dickel et al., 2021; Reid et al., 2021). Limited dispersal of individuals within a population can lead to genetic spatial autocorrelation reflected by an isolation by distance pattern (Aguillon et al., 2017). This pattern reflects the footprint left by drift and limited dispersal, but sometimes it can also result from spatial variation in environmental factors (Garroway et al., 2013) or non-random dispersal (Garant et al., 2005). External immigration can disrupt isolation by distance patterns by introducing genetic material from distant sources. Immigrants often exceed the number of locally born individuals in avian populations (Millon et al., 2019); they can impact local population dynamics and show phenotypic differences relative to local birds, affecting aspects like life-history traits and offspring dispersal behaviour (Quinn et al., 2011). Despite their importance, the state and characteristics of immigrants are typically unknown. One particular aspect that is not regularly taken into account is the relatedness among immigrants (Dickel et al., 2021). Common assumptions include immigrants being outbred and being unrelated to each other and locally born individuals. However, in a landscape where many small natural populations are connected by dispersal, this is not always true (Chen et al., 2016). An example of this is when there are reciprocal dispersal events between populations, whereby descendants of immigrants disperse back to their ancestors’ population. This situation is further aggravated by the fact that dispersal is not random and that more related individuals tend to have more similar dispersive behaviours (Matthysen et al., 2005). As the source of immigrants is often unknown in many studies, assuming they are unrelated and outbred may lead to inaccurate inferences.

Long-term data from a population of a small, short-lived passerine, the great tit (*Parus major*), provides an ideal system to explore local population structure and the effects of within-population dispersal and external immigration. This population is located in Wytham Woods, United Kingdom, it has been monitored since 1947 (Lack, 1964), and it represents a typical, large, continental population. Great tits in this population breed in nest boxes where, each spring, nestlings and adults are marked with unique metal leg rings. This information allows us to infer a social pedigree, which is often used as a proxy for kinship, although extra-pair paternity can bias relatedness estimates (Firth et al., 2015). An alternative approach that allows us to establish accurate genetic pedigrees is the use of genome-wide data, which together with dispersal data, can help us to understand the observed patterns of spatial genetic structure.

Here, we use two genomic datasets—one genotyped at ~10,000 SNPs from 2,644 individuals and another at ~600,000 SNPs from 949 individuals—to explore whether any population structure follows an isolation by distance pattern in autosomes and the Z chromosome separately and quantify how temporally persistent this pattern is. We then investigate the effects of immigration by i) exploring whether immigrants and locally born birds can be assigned to their own categories using a Random Forest classifier, ii) testing whether locally born great tits or immigrants are more related to each other, and iii) exploring the homozygosity levels of the locally born versus the immigrant pool, which can provide insights into the evolutionary potential of the population.

## Methods

### Study site and genotyping

Our study was conducted in Wytham Woods, a 385-hectare woodland close to Oxford, United Kingdom (51°46N, 1°20W). The site contains a little over 1200 nest boxes, of which 1019 are suitable for great tits, and 25-50% of them are occupied each spring by breeding great tits. All locally born birds born in nest boxes have been tagged with individual British Trust of Ornithology rings, and all birds caught as adults without a ring are considered to be immigrants, as nests in natural cavities are comparatively rare (Greenwood et al., 1979). They start breeding in their first year, and their average reproductive lifespan is 1.9 years.

A high-density SNP chip was designed based on sequencing of great tits from Europe and the United Kingdom (Laine et al., 2016; Kim et al., 2018). This dataset contains 949 samples of Wytham Woods great tits collected between 1983 and 2011, although most of the samples were collected between 2004-2009 (Table S1). The dataset contains probes for around 600,000 SNPs, of which 502,685 are polymorphic in our Wytham Wood samples.

For analyses requiring larger sample sizes and where the number of individuals is more critical than the number of SNPs, we used the 10,000-SNP chip developed by van Bers et al. (2014), which included 2,644 samples from Wytham birds (Table S1) and 4,725 polymorphic SNPs. The 600K dataset represents a subsample of the 10K, as 909 (95%) of the birds in the 600K are present in the 10K. Analyses based on the 600K and 10K datasets are summarised in Table S2. Great tits exhibit sex-biased dispersal, with locally born females dispersing approximately 1.5 times further than males—nearly 300 metres more—within Wytham Woods (Fig. S1). Given this pattern, gene flow is expected to show a different pattern on the sex chromosomes than on the autosomes (Ellegren, 2009). Therefore, we created two datasets, one in which we excluded the Z chromosome from our analyses as well as chromosome 1A since it contains a known large inversion that can bias downstream analyses (da Silva et al., 2019), and another one with the Z chromosome alone. For the 10K dataset, we retained only autosomal SNPs, as the Z chromosome did not contain enough markers. After filtering, the 600K autosomal dataset comprised 418,082 SNPs, the Z chromosome dataset contained 29,033 SNPs, and the 10K autosomal dataset included 4,070 SNPs.

Unless otherwise specified in the relevant sections, quality control was performed using PLINK (Purcell et al., 2007) with the following filtering criteria: genotyping frequency >95%, minor allele frequency (MAF) >0.05, and Hardy–Weinberg equilibrium P > 0.001, following Garroway et al., (2013).

### Population structure

Isolation by distance arises when genetic relatedness decreases as geographic distance increases. We first visualised the raw data by plotting the proportion of the genome that is identical by descent, representing genetic relatedness between two individuals and estimated with PLINK, against the distance between the breeding locations of the two individuals. To estimate identity by descent, which is sensitive to rare variants, we filtered out SNPs with MAF<0.1 (Browning & Browning, 2013). We expected identity by descent to decrease with distance. If this pattern was driven by transient spatial clustering of relatives, this effect would diminish as we excluded closer kin. To check this, we created plots including all relationships, then progressively removed first-degree, second-degree, and more distant relationships. SNP-based kinship degrees were calculated with the KING-robust kinship estimator through PLINK. We repeated this analysis by removing relationships inferred with the social pedigree (parent-offspring, full siblings and half siblings), and with natal distances rather than breeding distances.

We also visualised how patterns of isolation by distance differ between the autosomes and the Z chromosome. The Z chromosome spends two-thirds of its time in males and has a reduced effective population size, around three-quarters that of the autosomes (¾Ne) (Saunders & Muyle, 2024). This reduction in effective population size means that, all else being equal, we expect to see higher identity by descent on the Z chromosome compared to the autosomes (Cai et al., 2023). Additionally, because female great tits disperse further than males (Greenwood et al., 1979), we expect stronger isolation by distance among males than among females, across both autosomes and the Z chromosome.

We then formally tested for an isolation by distance pattern using the autosomes by building a Bayesian generalised additive mixed model in brms (Bürkner, 2017), incorporating: (i) identity by descent as the outcome variable, representing genetic relatedness between two individuals; (ii) the distance between breeding locations of the two individuals as an individual-level effect, modelled with a spline to allow for non-linear relationships, and different smooths for groups based on immigration status (both locally born, one locally born and one immigrant, or both immigrants); and (iii) a group-level effect for the year in which the birds were breeding. We added a multi-membership grouping term, to account for the fact that individuals are nested within more than one comparison. We used MCMC with four chains of 4000 iterations each, including a warm-up of 400 iterations. We evaluated convergence via visual inspection of the MCMC trace plots, checking that the ESS was greater than 200 and the R values for each parameter (R=1 at convergence).

### Renewal of the genetic basis of the isolation by distance pattern

The isolation by distance pattern is stable over time, but it reflects a dynamic process where the underlying genetic basis of isolation by distance is expected to change over time due to dispersal, mortality of close kin and external immigration. Therefore, we expect to see a decay in identity by descent within a certain area over time. To explore this, we compared identity by descent values between pairs of birds born in the same year and those born progressively further apart in time independently within the two most distant areas of the population. To choose these areas, we identified the two most distant nest boxes and selected 50 of the nearest nest boxes around each to represent the closest neighbours. We then estimated identity by descent between birds from these same areas across time. We also created a baseline that represents the average genetic relatedness between individuals from the most distant areas of the population in a given year, serving as a reference point.

### Identifying immigrants

Even if the linear combination of genome-wide SNP-based variation into orthogonal eigenvectors does not reveal single axes of variation that separate immigrants from residents, this is not incompatible with the existence of nonlinear combinations of those same eigenvectors that do. Our analysis used Random Forest classification on principal components derived from the genome-wide SNP data, following these steps using the SNPRelate (Zheng et al., 2012) and the randomForest (Liaw & Wiener, 2002) packages in R (R Core Team, 2013). First, to ensure balanced representation, we implemented a stratified sampling approach that equalised the number of samples from each population group and maintained this balance in both training (70%) and testing (30%) datasets, with undersampling of the majority class where necessary. We then applied dimensionality reduction through PCA to extract the most informative features from the highdimensional genomic data. The first 15 principal components, which captured 9.24% of the variance in the SNP data, were used as input features for our classification model. Using higher numbers of eigenvectors as input did not increase model performance.

We used a Random Forest algorithm to classify individuals as either immigrants or local birds based on these summary genomic features. To assess the robustness of our classification, we implemented a validation framework that included: a) multiple iterations of model training and testing with different random seeds to account for stochastic variation in sample selection and model operation; b) re-analysis with randomised labels to establish a baseline for classification performance; and c) out-of-bag error estimation to evaluate model accuracy during the forest-building process. We used multidimensional scaling (MDS) of the random forest proximity matrix to visualise the high-dimensional genomic similarities in a two-dimensional space. This gives a reasonable (if simplified) idea of the resulting model-based similarity space, and thus the degree to which immigrant and local birds are discernible. To further characterise the stability of our classification results, we conducted an ensemble analysis by running 50 random forests with different initialisations and averaging their proximity matrices and predictions. This produced a more robust representation of the genetic landscape and classification confidence for each individual.

### Genetic divergence across the genome between immigrants and local birds

By pooling all SNPs into a global PCA we might fail to detect differences between groups even if specific genomic regions of divergence, known as “islands of divergence”, exist. These regions may arise in the face of gene flow if local adaptation is ongoing (Nosil & Feder, 2012). To investigate this, we divided the dataset into immigrants and local birds and calculated *F*_*ST*_between these groups in 50 kb sliding windows using VCFtools (Danecek et al., 2011). This analysis was repeated using the top 50 most distinct immigrants based on MDS1 and the top 50 most distinct local birds based on MDS2, as identified from the Random Forest classification. We checked the kinship values of the two groups to confirm that these were not closely related individuals.

### Estimating homozygosity

We explored inbreeding by conducting Runs of Homozygosity (ROH) for each individual in PLINK. Two individual alleles can be homozygous by chance (“identical by state”), but contiguous lengths of homozygous genotypes that are present in an individual, also known as ROH, are a consequence of the parents transmitting identical haplotypes. The proportion of the genome in ROH can be used to measure inbreeding (Keller et al., 2011). We estimated ROH in PLINK using the autosomal dataset with no MAF threshold or LD pruning as recommended by Meyermans et al. (2020). We visualised the distributions of the average ROH length between immigrants and local birds.

## Results

### Spatial population structure

We conducted a genomic PCA on all breeding adults to visualise genetic variation within the population. The first two principal components (PC1 and PC2) explained 0.5% and 0.42% of the genetic variation, respectively (Fig. 1; Fig. S2). We found evidence of a fine-scale spatial genetic structure within the population, with a Northwest-Southeast cline represented by PC1. There was a negative correlation between identity by descent values and geographic distance, confirming an isolation by distance pattern (Fig. 1B). This pattern was most pronounced when close kin (parent-offspring or full-siblings) were included in the analysis, as expected when settlement probability decreases with distance. The isolation by distance effect diminished when close relatives were excluded (Fig. 1B). When we used the social pedigree, we found a similar pattern where exclusion of parent-offspring and full-siblings made the isolation by distance pattern weaker. However, excluding half-siblings did not have a big effect (Fig. S3).

**Fig. 1.**
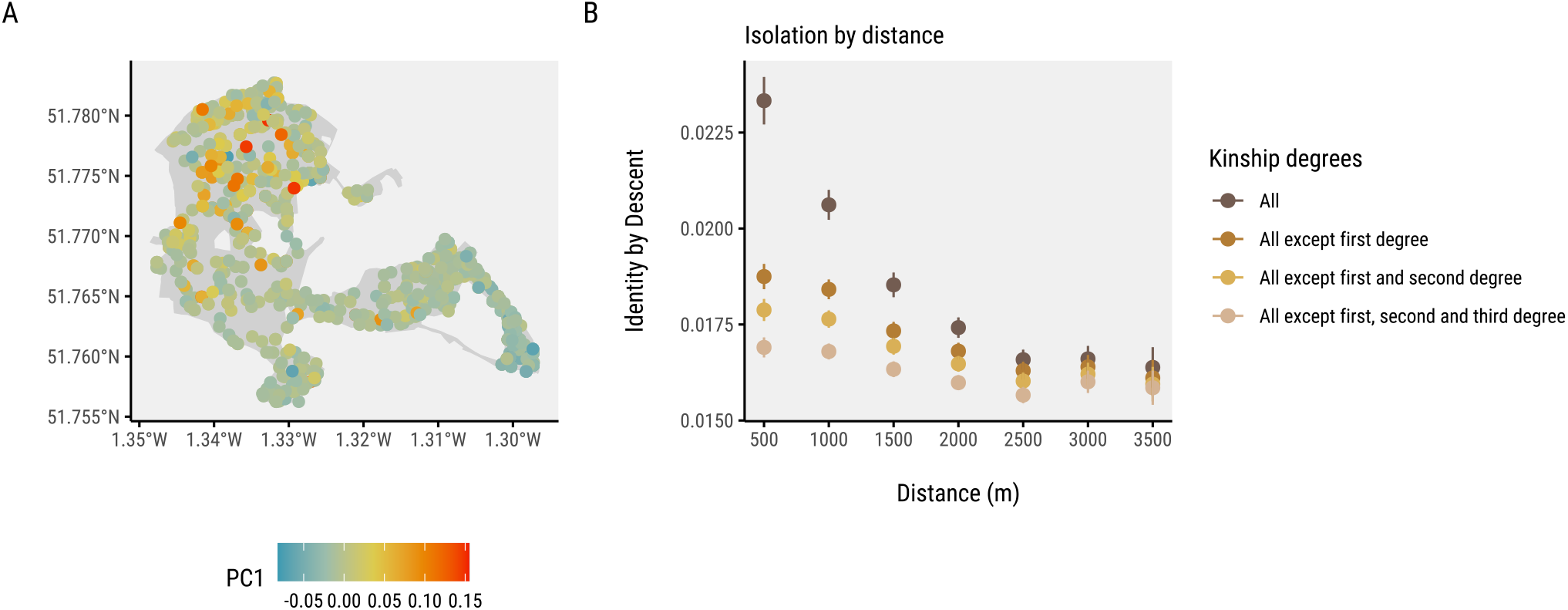
A) The genetic spatial structure of locally born breeding great tits in Wytham Woods is represented by PC1, the axis of variation that explains the most variance from a genomic PCA (0.5%). B) Identity-by-descent declines more sharply with breeding distance when close kin are included, but as these relatives are progressively excluded, the isolation by distance pattern weakens. This suggests that the observed genetic structure is largely driven by limited dispersal among closely related individuals.

We also compared isolation by distance using natal distance versus breeding distances (Fig. S4). Including all kinship degrees, natal distance revealed stronger genetic structure, with higher identity by descent at short distances and a steeper decline with increasing distance. However, this difference largely disappeared when first-degree relatives were excluded. As close relatives (first, second, and third degree) were progressively removed, the patterns for natal and breeding distances became increasingly similar, suggesting that the initial disparity was mainly due to the spatial clustering of close kin. When comparing the autosomes and the Z chromosome, we found a pattern of isolation by distance across all pair types on the autosomes, with the steepest decline in identity by descent in male-male pairwise comparisons (Fig. S5). Female-female and female-male comparisons also showed decreasing identity by descent with distance, but with gentler slopes and lower overall values. In contrast, on the Z chromosome, we only found a clear isolation by distance pattern in male-male pairs. Identity by descent values for female-female and female-male pairs were uniformly low across all distances.

Both locally born and immigrant individuals showed similar spatial genetic patterns, indicating a lack of strong differentiation between these groups (Fig. S6), and similar patterns of spatial population structure as immigrants (Fig. S2). We explored isolation by distance patterns among different pair types (locally born, immigrant, and mixed pairs). Locally born pairs showed a stronger isolation by distance pattern (*β*= −0.04; 95% CI: −0.06, −0.02) compared to locally born-immigrant pairs (*β*= −0.02; 95% CI: −0.03, 0.00), and immigrant-immigrant pairs (*β*= −0.01; 95% CI: −0.02, −0.01) (Fig. 2). Group-level effects showed no effect (*β*_multi-membershipterm_=0.1; 95% CI: −0.1, 0.1; *β*_year_=0; 95% CI: −0.1, 0.1) indicating consistent patterns across individuals and years. Locally born pairs showed higher identity by descent values when breeding at short distances than the other two categories, but lower values at long distances (Fig. 2).

**Fig. 2.**
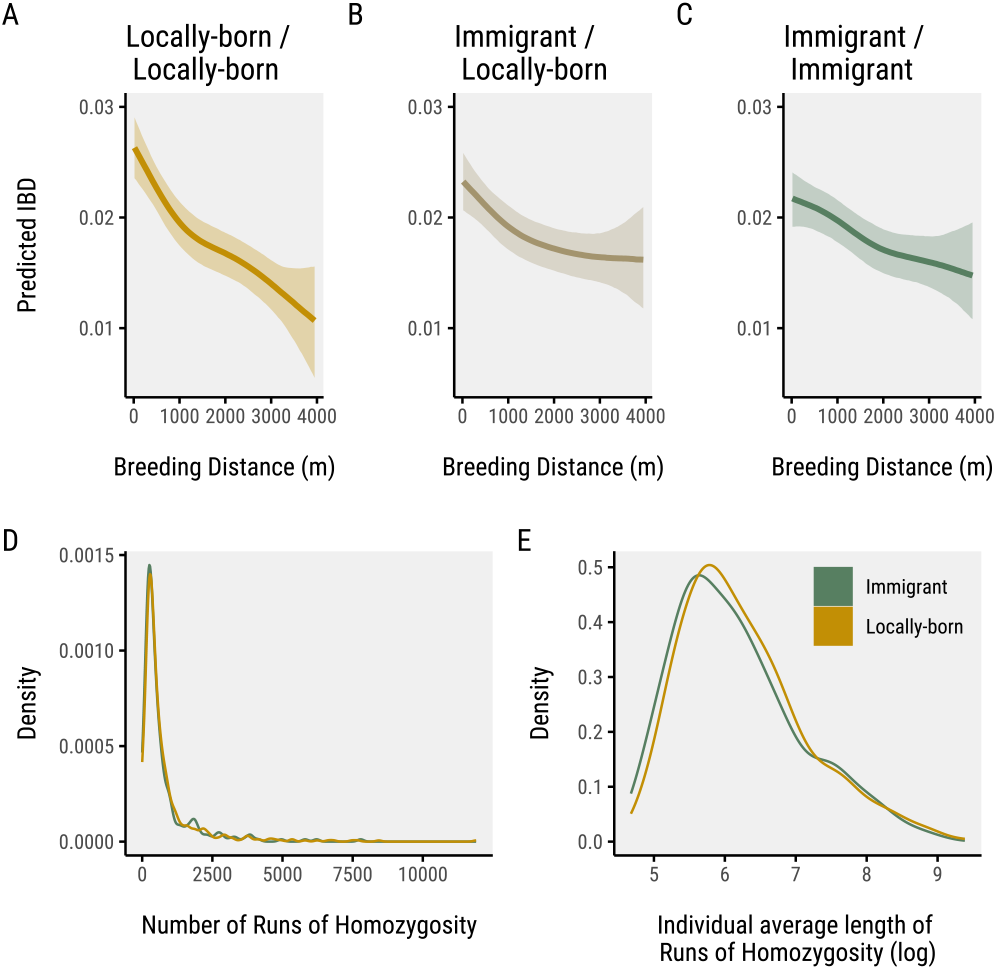
Posterior predictions of identity by descent with breeding distance indicate a stronger relationship for locally born pairs. This category also shows stronger identity by descent values than immigrants among each other at short distances.

Close kin relationships were rare across all pair types but showed clear differences in frequency depending on immigration status (Table S3). Locally born pairs had the highest proportion of close kin, with 0.0016 and 0.0031 being first- and second-degree relationships, respectively. In comparison, local–immigrant pairs showed lower proportions (0.0007 first-degree; 0.0015 second-degree), while immigrant–immigrant pairs had the fewest close relatives, with proportions of 0.0005 and 0.0012 for first- and second-degree kin, respectively. Despite these differences in kinship, both locally born and immigrant great tits exhibited similarly low levels of inbreeding, as reflected in comparable numbers and lengths of ROHs (Fig. 2).

### Renewal of the genetic basis of the isolation by distance pattern

We investigated the temporal dynamics of genetic structure by comparing identity by descent values between individuals born in the same year and those born increasingly far apart in time for two distinct and maximally separated parts of the population. The baseline estimate of identity by descent, indicating the average relatedness between birds from the most distant areas of the woodland, was 0.0016. In both sub-populations, we observed a temporal decline in identity by descent values, such that after 6–8 years, individuals from the same location were as genetically dissimilar as those separated by the full spatial extent of Wytham Woods (~ 4 km) within the same year (Fig. 3). This indicates that temporal separation of just a few generations can lead to levels of genetic differentiation comparable to those generated by several kilometres of spatial separation. In other words, being born 1 km apart in space has a similar effect on genetic relatedness as being born one generation apart in time. This corresponds well with the median natal dispersal distance in our population, which is also just slightly below 1 km. This pattern emerged in two independent areas, although in the northeastern corner (Fig. 3B) the decay was weaker than in the southeastern corner (Fig. 3C), and the initial identity by descent values were lower.

**Fig. 3.**
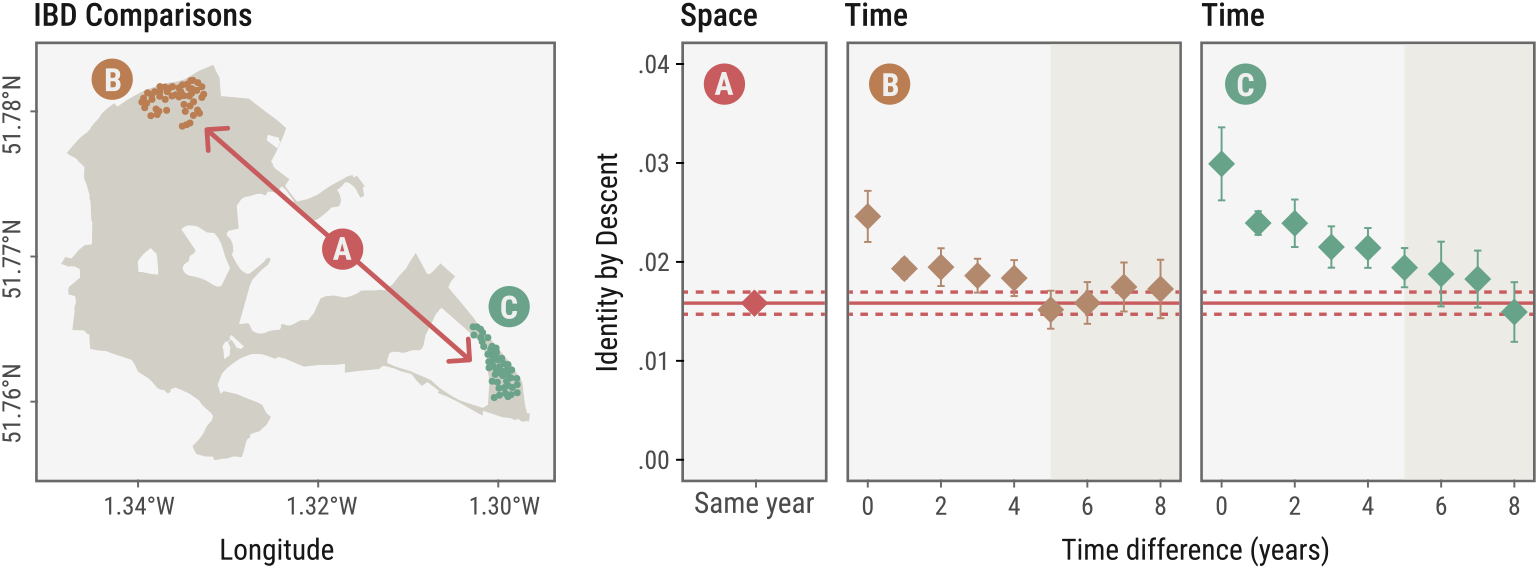
Birds show high identity by descent values with birds born in the same year and in the same area, but as time progresses, birds become less related due to population turnover (mortality, within-population dispersal and external immigration). Identity by descent values become indistinguishable from a baseline level, calculated as the average identity by descent between birds from the farthest sections of the woods within a given year, when comparing birds born 6 to 8 years apart.

### Identifying immigrants

The random forest classification based on genomic principal components achieved modest discriminatory power between immigrants and local birds (Fig. 4). The model had an overall accuracy of 64.8% (±2.7%), which, while statistically better than random chance, falls short of providing robust individual-level classification. The correct classification rate for locally born birds (69.0% ±3.1%) was somewhat higher than for immigrant birds (60.5% ±3.4%), suggesting greater genomic coherence among Wytham-born individuals. Our randomised (null) models performed as expected, with accuracy metrics clustered around 50% (balanced accuracy: 49.5% ±3.5%), confirming that the signal detected in this analysis, while modest, is genuine. These results suggest that while genomic differences between a subset of immigrant and locally born birds exist, they are subtle and insufficient for reliable individual assignment to either category, likely reflecting complex and overlapping patterns of genetic variation in the broader regional population.

**Fig. 4.**
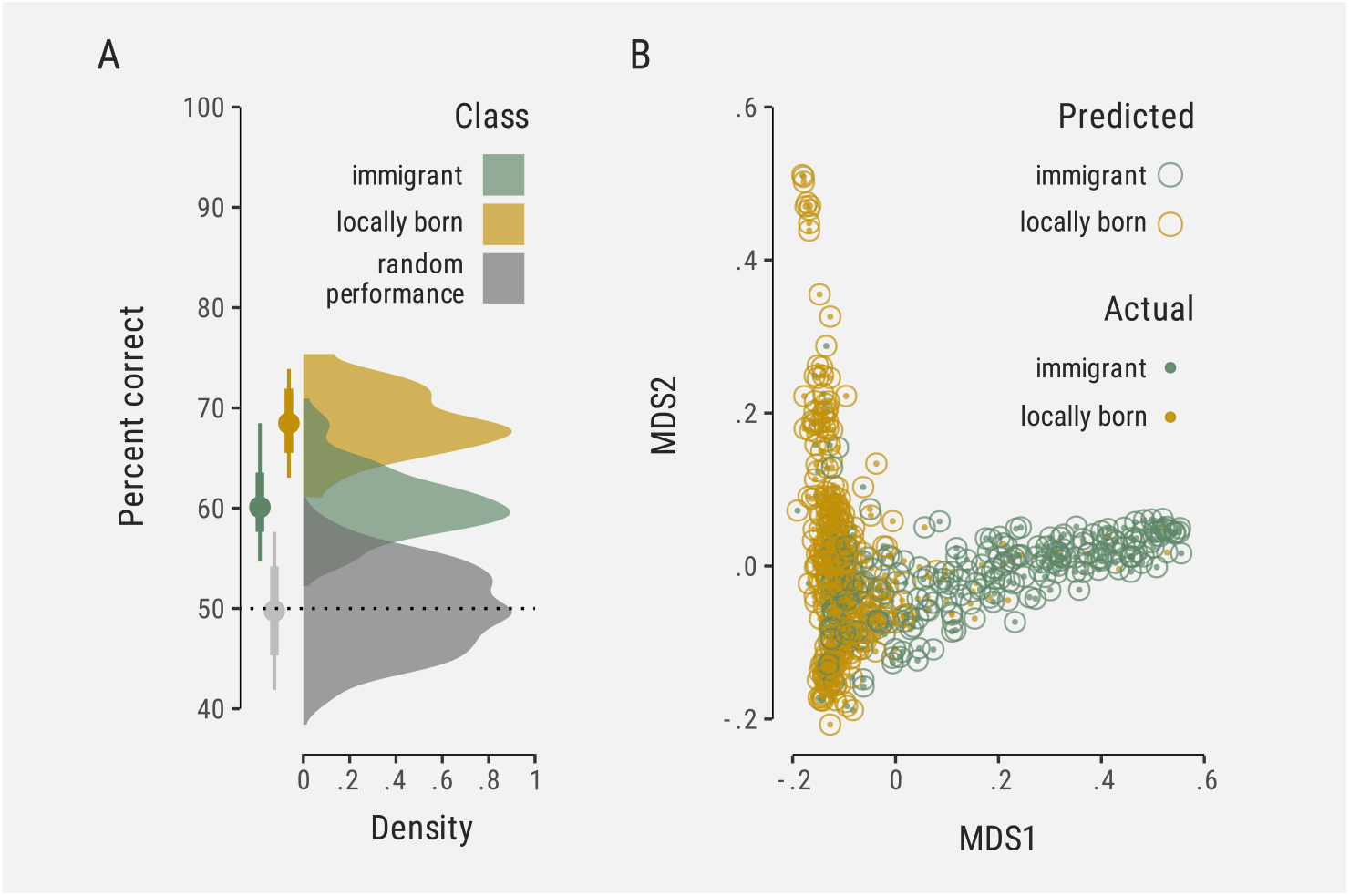
A random forest classifier trained on samples of locally born and immigrant birds achieves modest accuracy, with both groups, and especially locally born birds, partly distinguishable based on the main axes of genomic variation. A) Distribution of correct predictions in a binary classification task. B) First two dimensions in an MDS of the proximity matrix derived from the Random Forest classifier, showing how individual samples separate and overlap within this space.

### Genetic divergence across the genome between immigrants and local birds

When comparing all immigrants to locally born, we found an *F*_*ST*_ value of 3.64× 10^−4^, with similarly low *F*_*ST*_ values across the genome (Fig. S7). Comparing the top 50 immigrants and locally born identified by the Random Forest classifier inevitably showed a slightly higher *F*_*ST*_value of 1.05× 10^−3^, with higher windowed *F*_*ST*_values but no evidence of islands of divergence. Overall, *F*_*ST*_values were evenly distributed across the genome.

## Discussion

Using genomic data, we studied spatial genetic structure, immigration, and inbreeding in a long-term great tit population in Wytham Woods. We found: i) fine-scale isolation by distance driven by limited dispersal among close kin and ii) sex-biased dispersal, as reflected by different patterns in autosomes and the Z chromosomes; iii) the genetic basis underlying spatial structure is frequently renewed due to high population turnover; iv) modest genetic differences between immigrants and locals, with no distinct clusters; v) locally born birds are more genetically related to each other than to immigrants, which also show lower relatedness among themselves and, vi) similarly low inbreeding levels in both groups. These findings offer a clearer understanding of how dispersal shapes the genetic makeup of the Wytham Woods great tit population, and offer a perspective on how dispersal is likely to shape genetic structure in the many species that occupy, like this one, large continental meta-populations.

### Fine-scale spatial genetic structure reflects limited dispersal in close kin

We found a fine-scale spatial genetic structure within the population, which was further supported by a negative correlation between identity by descent values and geographic distance, indicating an isolation by distance pattern within the population as previously described by Garroway et al., (2013) with a less dense SNP chip and a different set of individuals. However, this pattern was most pronounced when closer kin were included in the analysis. As we removed degrees of kinship, the isolation by distance pattern diminished. This mostly results from closely related individuals, such as parent-offspring and full-siblings, remaining physically close together as breeders within neighbourhoods of contiguous territories. This interpretation is further supported by our comparison of natal and breeding distance metrics. When using natal distance, identity by descent values were substantially higher in the first bin (within 500 m) than when using breeding distances, which reinforces the idea that close kin are more likely to be found near each other at birth but tend to later breed at slightly greater distances. This is an inevitable observation in populations with dispersal distances that are smaller than their entire species distribution range, but there are extreme cases, such as in Florida scrub jays (*Aphelocoma coerulescens*) (Aguillon et al., 2017), where in only 10 km an isolation by distance pattern was evident and driven almost exclusively by limited dispersal (499 m). Our study uses a similar spatial scale and finds similar patterns, which aligns with what we would expect given the known dispersal range of great tits (Greenwood et al., 1979).

It is important to note that patterns of isolation by distance are not exclusively driven by limited dispersal; they can also arise from environmental heterogeneity or social factors, such as mating preferences or territory fidelity, that restrict genetic mixing across space. In this population, there is evidence of phenotype-dependent dispersal in a 40-year period (Garant et al., 2005) and associations between genetic variation and environmental factors that are themselves spatially variable, such as malaria infection risk and conspecific density (Garroway et al., 2013). Although we cannot exclude that nonrandom dispersal and environmental differences reinforce the spatial genetic variation we find, we see that limited dispersal of close kin explains much of the isolation by distance pattern.

### Sex-biased dispersal and its genomics consequences

In great tits, as in many bird species, females tend to disperse farther than males (Greenwood & Harvey, 1982; Michler et al., 2011). This pattern is generally attributed to differences in breeding roles: males benefit from remaining near their natal site due to their need to acquire and defend territories (Greenwood, 1980), while females gain more from dispersal by accessing a wider pool of potential mates. Female-biased dispersal may also help avoid inbreeding by reducing the chance of mating with close relatives. This behavioural asymmetry has clear genetic consequences. Because the Z chromosome spends most of its evolutionary time in males, restricted male movement leads to reduced gene flow on the Z and consequently stronger population structure relative to autosomes (Saunders & Muyle, 2024). Additionally, the Z chromosome has a smaller effective population size, making it more susceptible to drift and amplifying signals of relatedness such as identity by descent (Cai et al., 2023). These expectations align with our observed patterns: higher identity by descent values on the Z chromosome and male-male comparisons show stronger isolation by distance, which adds further support to conclude that limited dispersal is a key driver of spatial genetic structuring in this population. This is in line with theoretical expectations (Li & Kokko, 2019) and mirror results from other species with female sex-biased dispersal, such as sociable weavers (*Philetairus sociu*s) (van Dijk et al., 2015), Florida scrub jays (Aguillon et al., 2017), and male sex-biased dispersal in siberian jays (*Perisoreus infaustus*) (Li & Merilä, 2010), and white-browed sparrow-weaver (*Plocepasser mahali*) (Harrison et al., 2014). Together, our findings add to evidence that sex-specific dispersal influences relatedness and genetic structure differently on the sex chromosomes and autosomes.

### Temporal dynamics of genetic structure

We find that isolation by distance is consistent across years. However, the spatial genetic structure driving this pattern need not be static over time, as the population’s genetic landscape is dynamically shaped by two key processes: immigrants introducing new genetic variants into the population, and local loss of variants when individuals carrying them leave or die without reproducing. These demographic processes have been extensively explored in the population genetics literature, predicting that even in equilibrium, local extinction–recolonisation and dispersal can significantly alter genetic diversity and structure over time (Pannell & Charlesworth, 2000; Whitlock, 1992). Given the natal dispersal distance of Wytham great tits relative to the woodland’s extension, these results are qualitatively consistent with expectations driven exclusively by dispersal as described in stepping-stone models (Kimura & Weiss, 1964). However, this simplified framework—assuming random dispersal and no external immigration—does not capture the full complexity of spatial genetic structure in natural populations. Factors such as non-random dispersal (Garant et al., 2005), individual behavioural variation (Aplin et al., 2013), and each section’s geographic and demographic conditions all play important roles. For example, the slower and steadier decay of identity by descent observed in the southeastern corner of the study population (Fig. 3C) might reflect demographic effects of this being a relatively isolated patch, which might reduce the number of immigrants relative to emigrants, slowing population turnover and amplifying the effects of limited dispersal. In contrast, the northwestern corner (Fig. 3B) shows lower values of identity by descent among birds born the same year, which may reflect higher levels of immigration, leading to the faster “reshuffling” of alleles among subpopulations (Whitlock, 1992). Habitat structure and fragmentation are known to influence demographic and genetic turnover by altering how easily individuals can move between different parts of the environment. For example, in forest-dependent birds, even moderate habitat gaps can create significant barriers to dispersal, leading to detectable genetic structure across relatively short distances (Adams & Burg, 2015). In the context of Wytham Woods, variation in habitat connectivity, patch size, and edge effects may all contribute to differences in local immigration/emigration dynamics and their genetic consequences. While isolation by distance patterns may appear stable, they emerge from spatially variable and temporally dynamic demographic processes. Future studies could explicitly model how differential dispersal, influenced by factors such as geography, personality traits, and other constraints, shape the observed patterns of spatial genetic variation.

### Genomic evidence for limited divergence between locals and immigrants

Immigrants can increase genetic variation and reduce the probability of local adaptation (García-Navas et al., 2014), but this assumes they come from a separate genetic pool rather than being an extension or part of the local population. In our dataset, immigrants and local individuals do not form distinct clusters based on any single principal component or a simple combination of them. However, despite this overall genetic similarity, there are some detectable differences between the two groups. Using a Random Forest model with SNP data, we were able to classify locally born birds and immigrants at rates slightly above chance, though slightly less accurately for immigrants. This asymmetry might arise if locally born birds are more likely to intermix, creating recognisable genetic signatures, whereas immigrants, coming from various areas outside the population, share less genetic similarity (e.g. some birds labelled ‘immigrants’ are genetically very close to locally born birds, whereas others do not share recent ancestry with Wytham-born birds) and the model struggles to distinguish them as well. In fact, when we compared identity by descent across different pair types, we found that spatially proximate locally born birds were, on average, more related than immigrants, as well as having a higher proportion of close kin. This makes sense biologically, as immigrants are more likely to come from different areas outside of the woodland and will be less related, which aligns with the observations of Verhulst et al., (1997), who estimated that 94% of immigrants in Wytham Woods were born further than 2 km from the woodland. Additionally, immigrant pairs were more closely related at greater breeding distances than local pairs. This suggests that immigrants might disperse over longer distances, consistent with their lower isolation by distance pattern. However, we should consider sampling bias, as we only captured immigrants that successfully bred in the population, potentially selecting for individuals with higher dispersal and exploratory behaviours, a difference that Quinn et al., (2011) found between Wytham-born and immigrant birds.

Although genome-wide similarity suggests high gene flow, we next asked whether any islands of divergence exist between immigrants and local birds. Such divergence could reflect past or ongoing local adaptation, which can occur if either group experiences strong divergent selection, even in the presence of gene flow (Tigano & Friesen, 2016). We know that Wytham-born and immigrant birds differ in a few traits, including egg laying date (Verhulst et al., 1997), which is under selection (Garant et al., 2007), exploratory behaviour (Quinn et al., 2011), and song diversity (Merino Recalde et al., 2025). While some of these traits show low heritability (Jones et al., 2024; Quinn et al., 2009), islands of divergence may still be detectable, especially if the trait is underpinned by a simple genomic architecture or there is high linkage. To test this, we scanned the genome for differences between the two groups. However, we found negligible *F*_*ST*_values, with no clear outliers, which is consistent with our result that most immigrants are genetically very similar to locally born birds. We then examined a subset of individuals that were most distinguishable in the Random Forest analysis, and we observed a slight increase in *F*_*ST*_values across windows. However, these values remained negligible, and the outliers were spread evenly across the genome, with no obvious genomic islands of divergence. We consider that, at this spatial scale, this elevation in *F*_*ST*_ is more likely to be a result of neutral divergence rather than local adaptation of polygenic traits. This is because, in the face of gene flow, traits with simple genetic architectures and high linkage are more likely to persist and form genomic islands of divergence as they are less affected by recombination and more favoured by natural selection, maintaining local adaptation (Nosil & Feder, 2012; Tigano & Friesen, 2016). Therefore, the lack of genomic islands of divergence indicates that long-term local adaptation is unlikely to have taken place.

### Low inbreeding as a consequence of high dispersal and immigrant influx

Dispersal helps individuals to avoid inbreeding (Szulkin & Sheldon, 2008). In this population, the patterns of natal dispersal, combined with the fact that approximately 40% of the breeding population consists of immigrants annually, naturally reduce the chances of inbreeding. Our results indicate that birds have either no ROHs or a low number of short ROHs, which aligns with earlier findings that mating with close relatives is rare (1–2.6% of matings; Szulkin et al., 2007).

The high genetic diversity observed in great tits likely provides important advantages for population resilience in the face of environmental change. As great tits are known to be sensitive to phenological mismatches with their prey driven by climate change (Visser et al., 1998), maintaining genetic diversity may be crucial for adaptation to these shifting conditions. Immigration contributes to a high effective population size and introduces potentially beneficial genetic variants. Interestingly, high diversity occurs alongside the population’s capacity to develop local genetic structure, which can facilitate rapid adaptation to fine-scale environmental differences, especially when those conditions vary spatially. Together, these dynamics might represent an optimal balance: gene flow supports long-term evolutionary potential, while local genetic structure promotes short-term adaptive responses. This balance could be particularly critical as climate change continues to alter selective pressures in temperate woodland ecosystems.

## Conclusion

Our genomic analysis of the Wytham Woods great tit population reveals a fine-scale spatial genetic structure driven primarily by limited dispersal of close kin, with patterns that differ between autosomes and the Z chromosome in ways that reflect sex-biased dispersal. This genetic structure is temporally dynamic, with rapid turnover that likely constrains local adaptation. Most immigrants are genetically similar to local birds, though some are more distinct; local birds are generally more related to each other than to immigrants, which are also less related among themselves. Overall, the population remains genetically diverse, with enough gene flow to maintain evolutionary potential under changing environmental conditions.

## Acknowledgements

This work was supported by grants from the European Research Council (grants 250164 to BCS and 202487 to JS), Natural Environment Research Council (grant NE/J012599/1 to JS), and UKRI Frontiers award EP/X024520/1 to BCS.

## Author contributions

Conceptualisation: AE, BCS; Data Curation: AE, JS; Formal Analysis: AE (Bioinformatics), NMR (Random Forest); Funding Acquisition: BCS, JS; Investigation: AE; Methodology: AE, NMR; Visualization: AE, NMR; Writing - Original Draft: AE; Writing - Review & Editing: AE, NMR, JS, BSC.

### Data Availability Statement

Code to reproduce all analyses conducted in this paper can be found at https://github.com/andreaestandia/great-titgenomics and https://github.com/nilomr/greti-genomics. The SNP datasets were published by Kim et al., 2018 *Mol. Ecol. Resour*. and van Bers et al., 2012 *Mol. Ecol. Resour*.

