## Supplementary figures for "The genetic consequences of dispersal and immigration in a wild great tit population"

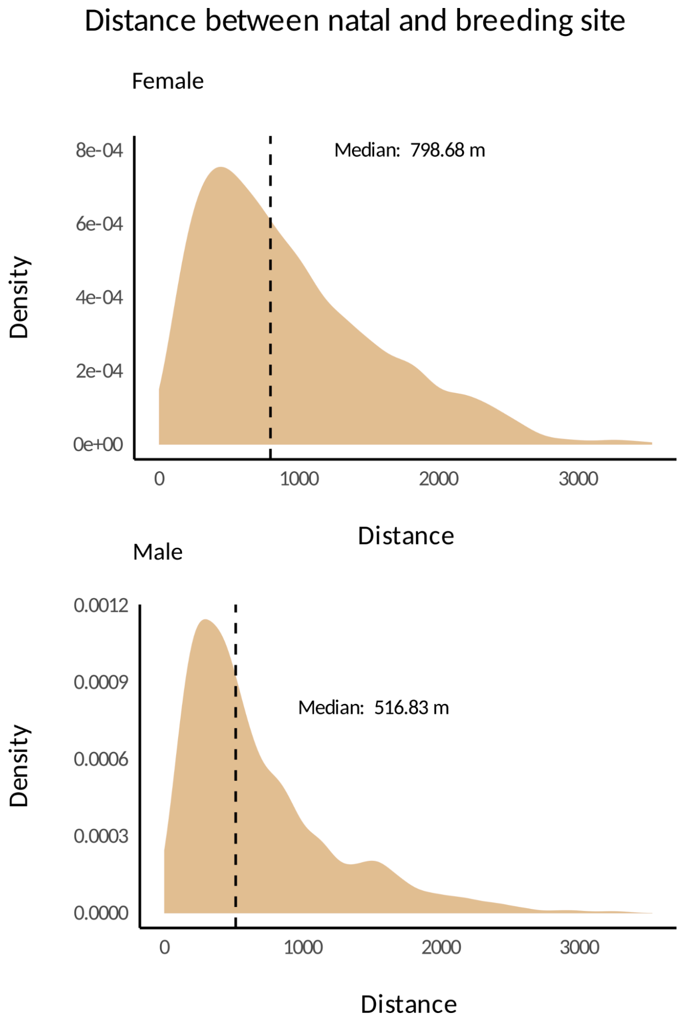


**Figure S1.** Sex-biased dispersal among locally-born individuals. Females dispersal almost 300m more than males do.

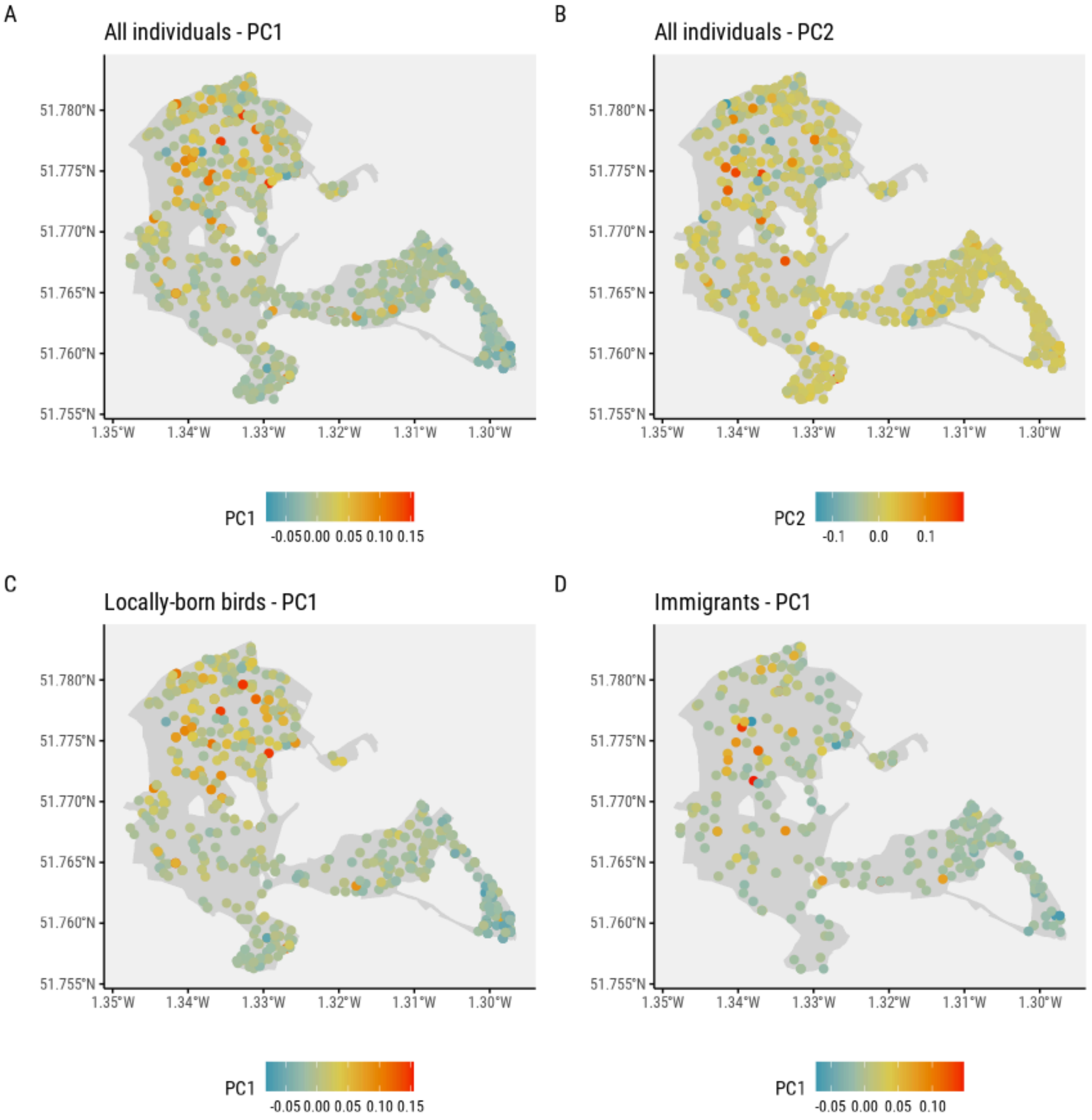


**Figure S2.** Axis of variation from a genomic PCA plotted onto a map of Wytham Woods in different subsets.

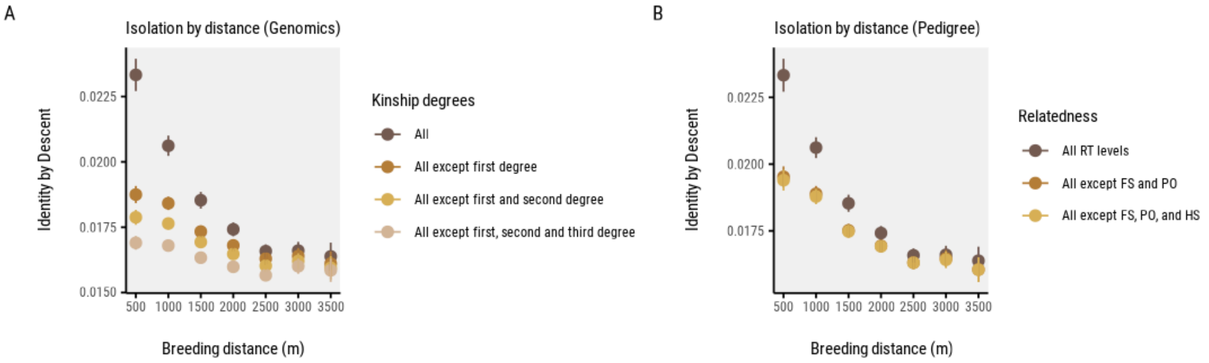


**Figure S3.** Isolation by distance by A) SNP-based kinship degrees and B) relatedness inferred using the social pedigree. RT=Relatedness, FS=Full siblings, PO=Parent-Offspring, HS=Half siblings.

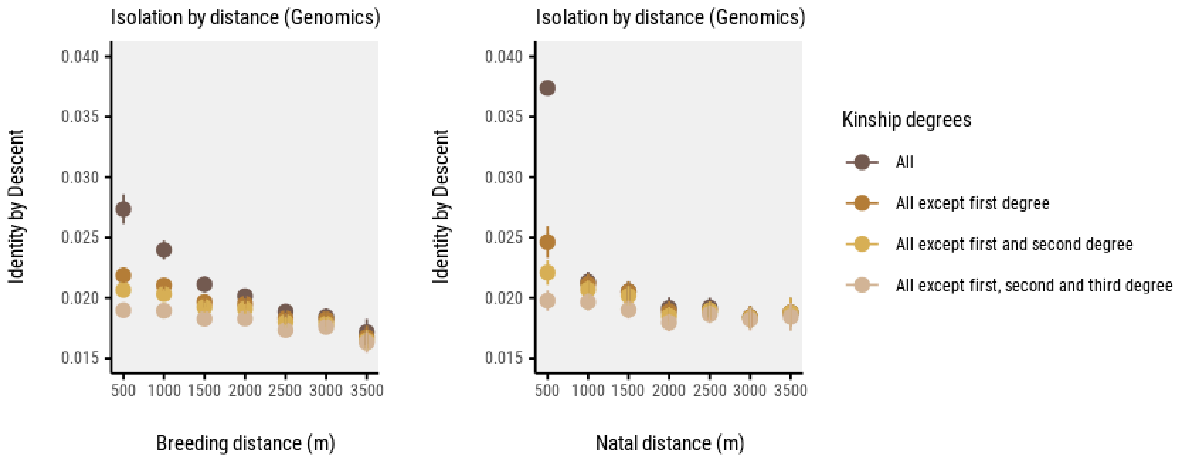


**Figure S4.** Isolation by distance using SNP-based kinship degrees with A) breeding distances and B) natal distances.

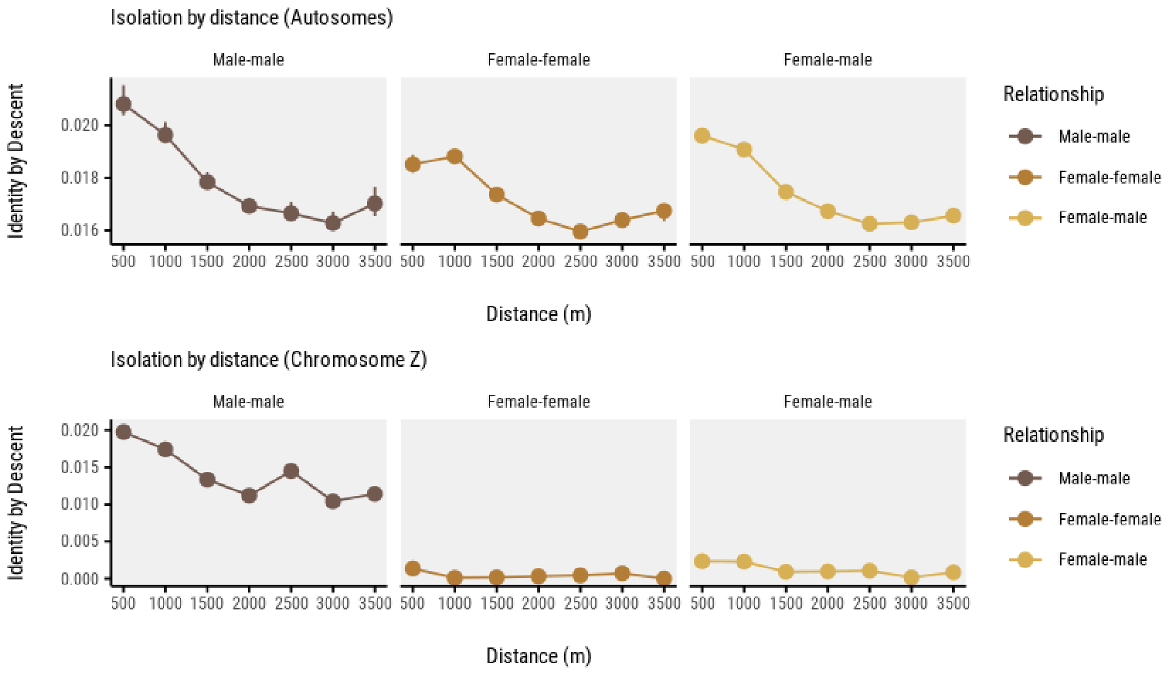


**Figure S5.** Genetic identity-by-descent across breeding distance for different sex-pair combinations on autosomes and the Z chromosome. Male-male pairs show consistently higher IBD values on both autosomes and the Z chromosome, particularly at shorter distances, suggesting sex-biased dispersal patterns. Female-female and female-male pairs exhibit similar declining IBD patterns with distance, though with generally lower values than male-male pairs, especially on the Z chromosome.

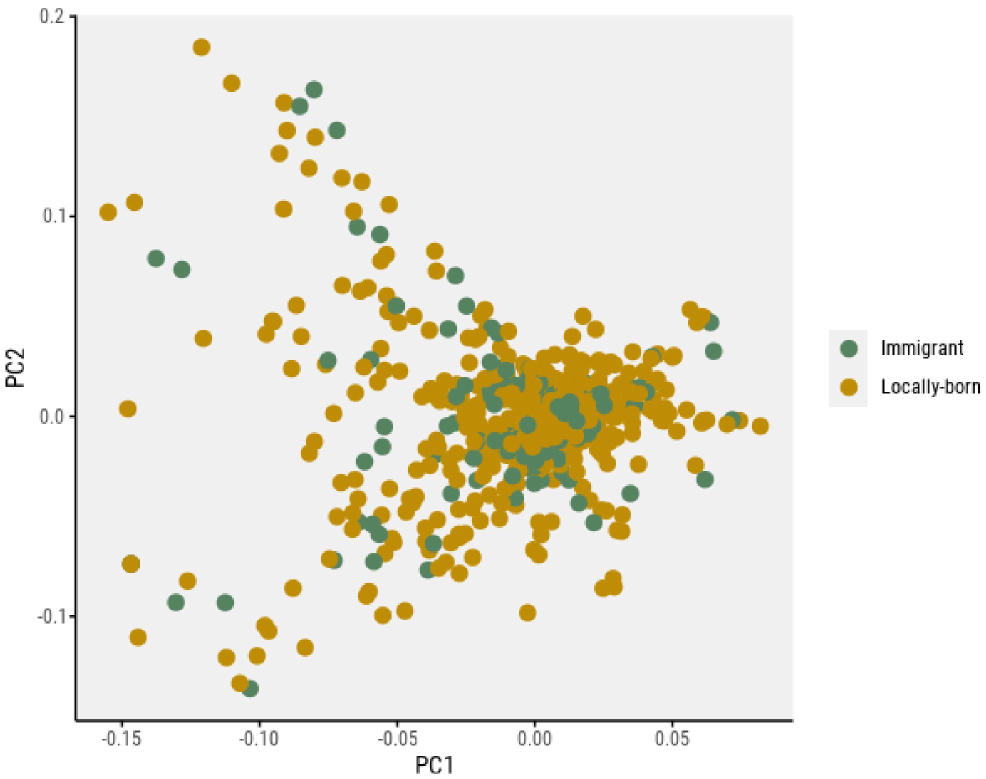


**Figure S6.** Locally-born and immigrant birds are not clustered in distinct genetic groups when doing a genomic PCA.

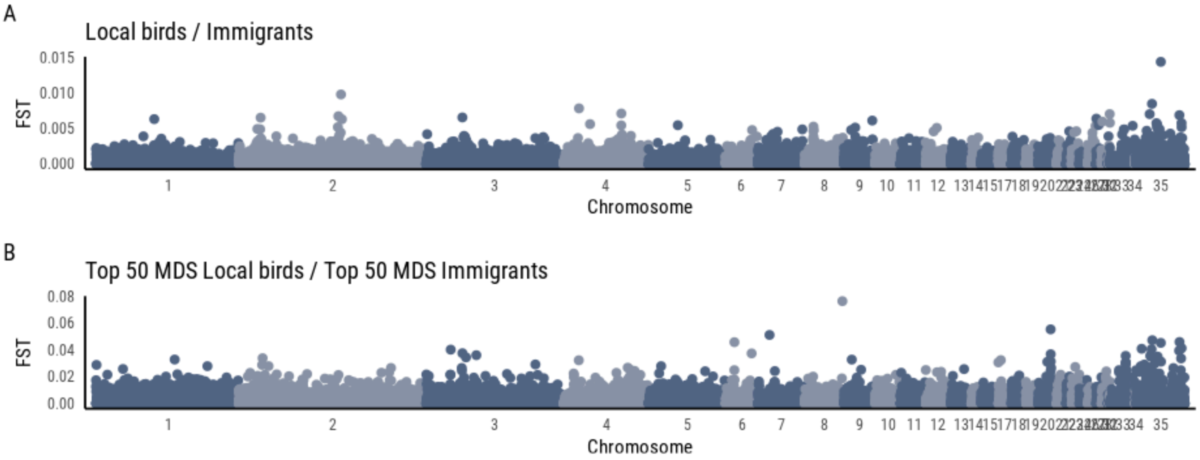


**Figure S7.** Fst was calculated in 50 kb windows across the genome, comparing A) all locally born birds and immigrants, and B) the top 50 most distinct immigrants based on MDS1 and the top 50 most distinct local birds based on MDS2, as identified from the Random Forest classification.

### Supplementary Tables captions

**Table S1.** Sample information for great tit individuals genotyped at 10K and 600K SNPs from the Wytham Woods population. Each row represents a unique individual identified by its BTO ring. Columns include the year of birth, nest box location, dataset, parental BTO ring codes (mother and father), sex (if known), and immigration status (resident or immigrant).

**Table S2.** Type of dataset used for each analysis as different analyses require a balance between the number of individuals and SNP density. For analyses like PCA, which help detect clines in population structure, or for those that model kinship, we use the 10K dataset because it includes more individuals. In contrast, analyses that require higher SNP density, such as window-based Fst calculations, are performed with the 600K dataset. Note that the 600K dataset is a subset of individuals from the 10K dataset.

**Table S3.** Pairwise kinship classification by immigration status in the Wytham Woods great tit population. Pairs are categorised by type (Immigrant/Immigrant, Resident/Immigrant, or Resident/Resident) and kinship degree (First degree, Second degree, or Unrelated). The table reports the number of pairs (N) and the proportion each kinship category represents within its pair type.
